# Neural stem cell interkinetic nuclear migration is controlled by a phosphatidylinositol transfer protein/non-canonical planar cell polarity signaling axis

**DOI:** 10.1101/2020.12.17.423231

**Authors:** Zhigang Xie, Vytas A. Bankaitis

## Abstract

The mammalian neocortex undergoes explosive expansion during embryonic development. From an evolutionary perspective, higher complexity of the neocortex is accompanied by a prominent expansion in its lateral dimension so that the neocortical surface area is increased. Expansion in the radial dimension throughout evolution is limited so that neocortical thickness is strongly restricted^1–3^. The underlying mechanisms for restricting neocortical thickness remain unclear. Expansion of the developing mouse neocortex is driven by neurogenesis which is itself primarily fueled by neural stem cells (NSCs). NSCs form a pseudostratified epithelium and exhibit a hallmark cell cycle-dependent nuclear movement termed interkinetic nuclear migration (IKNM) ^2–4^. While IKNM plays a critical role in cell fate determination, it remains a poorly understood process. Herein, we demonstrate IKNM relies on a phosphatidylinositol transfer protein (PITP)-noncanonical planar cell polarity (ncPCP) signaling axis that restricts radial expansion of the developing neocortex. Ablation of PITPα/PITPβ in NSCs compromised IKNM -- resulting in a thickened neocortex and perturbed curvature of its ventricular surface. Those phenotypic derangements in IKNM and neocortical morphogenesis were recapitulated in mouse embryos individually ablated for two ncPCP receptor gene activities and in a mosaic neocortex expressing a dominant-negative variant of a third ncPCP receptor. Finally, PITP signaling links to ncPCP pathway activity by promoting membrane trafficking of a subset of ncPCP receptors from the trans-Golgi network to the NSC cell surface. We conclude IKNM is a driving force for a special form of convergent extension regulated by coupling PITP-mediated phosphoinositide signaling with activity of the evolutionarily conserved ncPCP pathway.

---

PITPs are evolutionarily conserved proteins that fall into two distinct classes typified by the Sec14- and StAR-related lipid transfer (StART) structural folds^5^. Although these two PITP classes are structurally unrelated, PITPs of both execute similar functions in cells by potentiating phosphoinositide metabolism and signaling^5^. In particular, PITPs stimulate production of phosphatidylinositol-4-phosphate (PtdIns-4-P) by enhancing the activities of PtdIns 4-OH kinases^5^. Mammalian PITPNA and PITPNB are highly homologous StART-like PITPs that are expressed in NSCs of embryonic mammalian brain and function cooperatively to support development of the mammalian neocortex^6^. Ablation of both PITP isoforms in NSCs results in a catastrophic failure in neocortical development signified by cell-autonomous loss of NSC radial polarity followed by a rampant apoptosis that eliminates the developing dorsal forebrain. From the cellular perspective, PITPNA/PITPNB are required for production of a PtdIns-4-P pool on trans-Golgi network (TGN)/endosomal membranes that recruits the nonconventional myosin (MYO-18A)-binding protein GOLPH3 to those compartments. This recruitment in turn chaperones the asymmetric MYO18A- and F-actin directed loading of the TGN network into the apical processes of NSCs^6^. However, the mechanisms by which PITP-dependent PtdIns-4-P signaling in TGN/endosomal membranes interfaces with NSC biology and neocortical development remain obscure. Herein, we demonstrate PITP-dependent signaling directly interfaces with signaling via the non-canonical planar cell polarity (ncPCP) pathway to regulate NSC IKNM in support of a convergent extension mechanism that regulates morphogenesis of the developing mammalian neocortex.

## PITP-deficiencies result in neocortex dysmorphologies and impair IKNM

An *Emx1-Cre* driver^7^ was used to induce dorsal forebrain-specific deletion of the *Pitpna* and *Pitpnb* genes. Widespread apoptosis was observed in PITP double knockout (DKO) neocortices (*Pitpna^fl/fl^ Pitpnb^fl/fl^ Emx1^Cre/+ or^ ^Cre/Cre^*) at E13.5 and later stages^6^. At earlier stages, however, apoptosis was sparse^6^, permitting analysis of interpretable phenotypes at the tissue level. In that regard, the medial region of PITP DKO neocortices were notably thicker and shorter in the mediolateral dimension when compared to controls at early E12 (Extended Data Fig. 1a, b). This phenotype not only became more exaggerated by late E12 (Fig. 1a, b), but was accompanied an obvious reversal in ventricular surface curvature from concave in control neocortices to convex configurations in PITP DKO neocortices (Fig. 1a, b; arrows). This transformation from concave to convex curvature was consistent with increased numbers of cell bodies at the ventricular surface of PITP DKO neocortices relative to the numbers of cell bodies at the VZ/subventricular zone border. These dysmorphologies were clearly evident despite the fact that cells lining the ventricular surface were predominantly NSCs -- as was the case for the wild-type condition (Extended Data Fig. 1c, d). Moreover, NSC mitoses appropriately occurred mostly along the ventricular surface^6^ (Fig. 1c, d). These data suggested that reconfiguration of the ventricular surface from a concave to a convex curvature resulted from IKNM defects where nuclei of newborn NSCs were unable to efficiently migrate away from the ventricular surface.

**Fig. 1.**
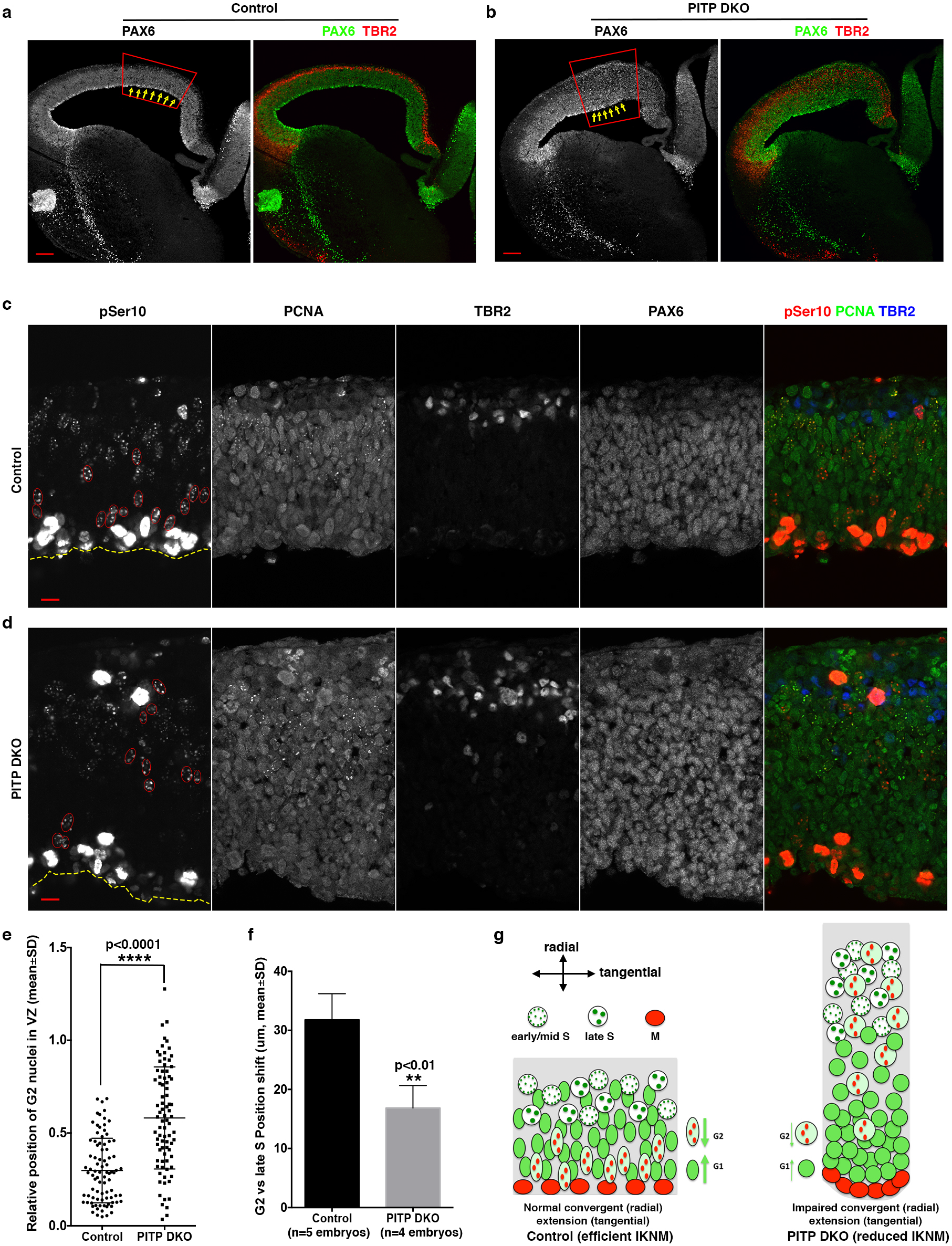
Altered neocortical shape and impaired IKNM in PITP DKO embryos. **a, b,** Markedly increased thickness (boxed areas) and convex ventricular surface (indicated by arrows in **b,** compared to the area indicated by arrows in **a**) were observed in the medial part of the neocortex in PITP DKO embryos at late E12. Images are representative of at least 4 embryos for each group. Scale bars: 100μm. **c-f,** IKNM defects in PITP DKO mutant embryos at E12.5. Representative images are shown in **c** and **d**. Quantifications were performed on the medial part of the neocortex in embryos at the stage of E12.5 (regardless of whether the stage was early E12 or late E12). G2 nuclei (large puncta of pSer10 and diffuse nuclear PCNA) are outlined in red, and the ventricular surface is outlined by dashed yellow lines. Scale bars: 10μm. **e**, The relative positions of G2 phase NSC nuclei were significantly shifted between control and PITP DKO embryos. Quantification represents G2 phase NSC data sets pooled from at least 4 independent embryos for each group. **f,** Shift in nuclear position toward the ventricular surface in NSCs transitioning from late S to G2 phase was significantly reduced in PITP DKO embryos compared to control. **p<0.01, ****p<0.0001, Student’s t-test. **g,** Convergent extension model of IKNM. This model rationalizes the neocortical shape changes (increased thickness and convex ventricular surface) in PITP DKO embryos.

To determine whether IKNM was compromised in PITP DKO neocortices, a reliable method for distinguishing NSCs at different stages of the cell cycle was required. Thus, the positions of NSC nuclei (i.e. nuclei of PAX6^+^TBR2^−^ cells) were monitored at different phases of the cell cycle by co-immunostaining for PCNA and Ser10-phosphorylated histone H3 (pSer10). PCNA is expressed in all proliferating cells but exhibits signature cell cycle-dependent nuclear distribution patterns. PCNA exhibits a diffuse nuclear localization in G1 and G2 cells but rearranges to a punctate localization profile during S-phase^8–10^. Whereas pSer10 is commonly used as a marker for mitotic cells, it also marks G2 -phase cells^11^. Interestingly, pSer10 co-localized with the large majority of PCNA puncta and its immunostaining intensity varied significantly as a function of the cell cycle. Signal intensity was extremely weak in NSCs with small nuclear PCNA puncta (i.e. in early and mid S-phase cells), was modestly increased in NSCs with large nuclear PCNA puncta (late S-phase cells), was further intensified in NSCs with pSer10^+^ nuclear puncta and diffuse nuclear PCNA, and was dramatically intensified in NSCs with pSer10-labeled chromatin (M-phase cells; Extended Data Fig. 2a). These profiles, when coupled with demonstrations that both PCNA and pSer10 label centromeric regions^11, 12^, and that PCNA distributions transformed from punctate to diffuse profiles upon entry into G2^8–10, 12^, identified NSCs with large nuclear pSer10 puncta and diffuse nuclear PCNA as G2-phase cells (Extended Data Fig. 2b). Moreover, NSCs marked by punctate nuclear PCNA were binned as S-phase cells regardless of pSer10 profile, whereas NSCs with diffuse nuclear PCNA staining and no detectable pSer10 fluorescence were classified as G1-phase cells (Extended Data Fig. 2b). By these criteria, late S- and G2-phase NSCs were unambiguously identified. Finally, G1- and early S-phase NSCs were confidently recognized in the apical quartile of the ventricular zone (VZ) as this is a region where mid to late S-phase cells were rare.

Late S-phase NSC nuclei (PAX6^+^TBR2^−^) were primarily distributed in the basal half of the VZ in control neocortices (Extended Data Fig. 2c, 3a), G2-phase NSC nuclei were predominantly distributed throughout the apical half of the VZ (Extended Data fig. 2c; Fig. 1c, e), and M-phase NSC nuclei lined the ventricular (apical) surface (Fig. 1c). Those observations are consistent with previous IKNM models in which M-phase nuclei localize to the ventricular surface of the neocortex, migrate away in a basally-directed fashion to the upper VZ of the neocortex during G1, complete S-phase in the basal side of the VZ, and subsequently retreat back toward the NSC ventricular surface during G2.

Three observations diagnosed impaired IKNM in PITP DKO neocortices. First, whereas late S-phase DKO NSC nuclei remained distributed predominantly in the basal half of the VZ (Extended Data Fig. 2c, 3a), G2-phase NSC nuclei failed to normally populate the apical half of the VZ (Fig. 1d, e). Second, the relocalization of NSC nuclei toward the ventricular surface during the late S-to G2-transition was significantly diminished (Fig. 1f). Third, the fractional representation of G1/early S-phase NSC nuclei was significantly elevated in the ventricular quartile of the VZ (Extended Data Fig. 3b). In aggregate, these data indicate that PITP-deficiencies: (i) impair apically-directed migration of G2-phase NSC nuclei toward the ventricular surface, and (ii) impair basally-directed migration of G1-phase NSC nuclei toward the VZ/subventricular zone boundary.

## IKNM and convergent extension of the developing neocortex

As morphology of the murine E12.5 neocortex is primarily contoured by the VZ, impaired IKNM suggested a rationale for the observed derangements in neocortical morphology. This rationale was conceptualized as promoting convergent extension of the developing neocortex. Given that mitosis occurs at the ventricular surface, the data are consistent with a mechanism where IKNM provides a force that exerts two effects. First, IKNM entices the VZ to converge radially as the constitutive migration of G1 and G2 nuclei toward each other contracts the VZ in the radial dimension. Second, IKNM promotes lateral extension of the VZ as migrating NSC cell bodies crowd neighboring cell bodies (Fig. 1g). Considering an extreme case, a complete IKNM block would channel VZ expansion in the radial dimension by continuously adding cell bodies of newly born NSCs to the ventricular surface. The consequences of such a block would be: (i) a thicker VZ due to enhanced radial expansion, (ii) a VZ severely restricted in the mediolateral dimension as lateral expansion is inhibited, and (iii) a convex ventricular surface deformed by excessive accumulation of cell bodies at the ventricular surface relative to the strongly reduced numbers of cell bodies in outer layers of developing neocortex. Accordingly, partial inhibition of IKNM would result in proportionately graded derangements in VZ thickness, VZ mediolateral length and curvature of the ventricular surface. This concept formalizes IKNM as a significant driving force for a specialized form of convergent extension -- a tissue morphogenetic process employed in multiple developmental contexts^13, 14^. Several independent approaches were undertaken to test this hypothesis.

## Non-canonical Wnt/PCP receptor-deficiencies phenocopy the neocortical dysmorphologies and impaired IKNM of PITP-deficient NSCs

One direct prediction of a convergent extension model for IKNM is that the non-canonical ncPCP pathway, the evolutionarily conserved core signaling circuit for convergent extension^13, 14^, is required for IKNM and neocortical morphogenesis. This prediction was interrogated using several independent experimental approaches that examined whether specific ablation of the ncPCP pathway recapitulated the neocortical and IKNM derangements observed for PITP DKO embryos. In that regard, several receptors function specifically in promoting ncPCP signaling (i.e. VANGL1/2, CELSR1/2/3, PTK7)^15, 16^ and, importantly, are not involved in signaling via the canonical Wnt/β-catenin pathway. These specificities permitted clean distinctions between functional involvements of the ncPCP vs the canonical Wnt/β-catenin pathway.

Examination of *Vangl2^lp/lp^* embryos at E12.5 revealed the mutant embryos exhibited severe anatomical defects in the central nervous system. These included failures in neural tube closure (as previously reported)^17, 18^ and rupture of the neocortex (Extended Data Fig. 4a). While these defects might indirectly affect neocortical morphogenesis, we noted that increased thickness of the neocortex and a convex ventricular surface were common features in intact neocortical areas physically distant from the site of rupture. These features were particularly striking in the medial neocortex (Fig. 2a, b). Alterations in neocortical length in the mediolateral dimension were not assessed because the significant derangement of overall forebrain morphology in the mutant embryos precluded accurate comparisons of that parameter between mutant and control embryos. However, IKNM defects were obvious in those thickened regions of the neocortex as indicated by: (i) altered spatial distributions of G2 NSC nuclei but not of late S-phase NSC nuclei (Fig. 2c-e; Extended Data Fig. 4b), (ii) marked reduction in the apically-directed migration of G2-phase nuclei toward the ventricular surface (Fig. 2f), and (iii) pronounced accumulation of G1/early S-phase NSC nuclei in the apical quartile of the VZ (Extended Data Fig. 4c).

**Fig. 2.**
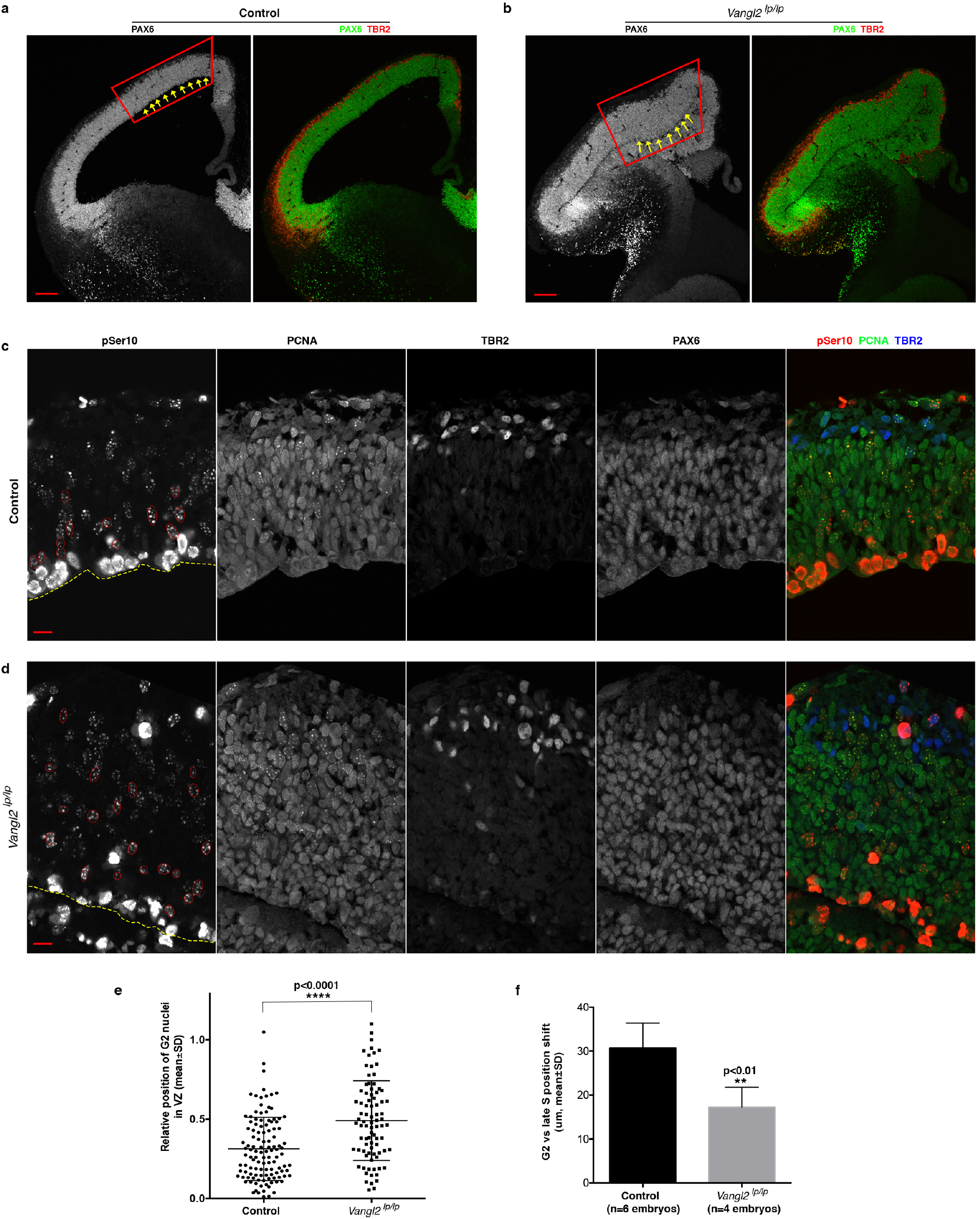
Altered neocortical shape and impaired IKNM in *Vangl2^lp/lp^* mutant embryos. **a, b,** Increased neocortical thickness (boxed areas) and convex ventricular surface (indicated by arrows in **b,** compared to the area indicated by arrows in **a**) were common in *Vangl2^lp/lp^* mutant embryos at E12.5, particularly in the medial neocortex. Images are representative of at least 5 embryos for each group. Scale bars: 100μm. **c-f,** IKNM defects in *Vangl2^lp/lp^* mutant embryos at E12.5. Representative images are shown in **c** and **d**. Neocortical areas with increased thickness were examined for IKNM defects. G2 nuclei (large puncta of pSer10 and diffuse nuclear PCNA) are outlined in red, and the ventricular surface is outlined by dashed yellow lines. Scale bars: 10μm. **e**, The relative position of G2 phase NSC nuclei was significantly different between control and *Vangl2^lp/lp^* mutant embryos. G2 phase NSCs pooled from 4 or more embryos for each group were used for quantification. **f,** The nuclear position shift toward the ventricular surface in NSCs progressing from late S to G2 phase was significantly reduced in *Vangl2^lp/lp^* mutant embryos compared to control. **p<0.01, ****p<0.0001, Student’s t-test.

Embryos homozygous for the *curly tail bobber (ctb)* allele of the ncPCP receptor *Celsr1* were similarly analyzed. This mutant mouse carries a single nucleotide deletion in Exon 5 of the *Celsr1* gene, resulting in a null allele (c.4614delC; p.Ser1538RfsX35) where a translational frame-shift induces downstream nonsense-mediated mRNA decay^19^. Consistent with previous reports of incomplete penetrance of tissue defects in *Celsr1* null mice^20, 21^, 53% of *Celsr1^ctb/ctb^* embryos (18/34) presented open neural tube defects. Because the lack of obvious neural tube defects in some nullizygous embryos likely reflected active compensatory mechanisms^18^, analyses were restricted to mutant embryos with neural tube defects. As observed for *Vangl2^lp/lp^* embryos, *Celsr1^ctb/ctb^* homozygotes exhibited neocortical shape changes and IKNM defects. Moreover, these were phenotypically indistinguishable from those recorded for the *Vangl2^lp/lp^* embryos. That is, the cardinal phenotypes included: (i) rupture of the neocortex (Extended Data Fig. 5), (ii) a thickening of the medial neocortex accompanied by a convex ventricular surface (Extended Data Fig. 6a, b), (iii) reduced distribution of G2, but not late S-phase, NSC nuclei in the apical half of the VZ (Extended Data Fig. 6c-f), (iv) decreased shift of nuclear position toward the ventricular surface during NSC transition from late S-to the G2-phase of the cell cycle (Extended Data Fig. 6g), and accumulation of G1/early S-phase NSC nuclei in the apical quartile of the VZ (Extended Data Fig. 6h). Thus, both *Vangl2^lp/lp^* and *Celsr1^ctb/ctb^* mutant embryos not only shared similar defects in neocortical morphogenesis and IKNM consistent with a convergent extension model for IKNM, but these also phenocopied the defects recognized in PITP DKO neocortex.

## Recapitulation of the neocortical dysmorphologies and impaired IKNM of PITP-deficient NSCs in a mosaic model for ncPCP deficiency

The *Vangl2l^p/lp^* and *Celsr1^ctb/ctb^* data were further buttressed by an independent experiment that addressed whether functional interference with a third receptor specific to the PCP pathway (PTK7) phenocopied the defects in neocortical morphogenesis and IKNM scored for PITP DKO, *Vangl2l^p/lp^* and *Celsr1^ctb/ctb^* mutant embryos. To overcome concerns that interpretation of mutant neocortices can potentially be confounded by indirect effects, we adopted an experimental regime that did not rely on analyses of mutant mice. Rather, a regional mosaicism was induced in an otherwise wild-type neocortex by introducing a dominant-negative allele of *Ptk7* into the neocortex of wild-type E12.5 mouse embryos by in utero electroporation. This dominant-negative *Ptk7* allele encodes a truncated form of the receptor that lacks the cytoplasmic tail (PTK7ΔC)^22, 23^. Morphological status of the neocortex and IKNM activity were subsequently assessed after 24h. As a high fractional representation of electroporated NSCs is required to influence neocortex morphology, only embryos with high transfection efficiencies were analyzed. Indeed, PTK7ΔC expression induced local deformation of the normally concave ventricular surface to a convex contour and evoked increased tissue thickness exclusively in efficiently transfected neocortical regions (Fig. 3a, b). Moreover, PCNA/pSer10 co-immunostaining demonstrated those abnormally thickened regions of the neocortex exhibited impaired IKNM (Fig. 3c-f; Extended Data Fig. 7). Thus, interfering with ncPCP signaling in a restricted region of an otherwise wild-type neocortex was sufficient to evoke local neocortical dysmorphologies and IKNM defects that phenocopied those recorded for PITP DKO, *Vangl2l^p/lp^* and *Celsr1^ctb/ctb^* embryos.

**Fig. 3.**
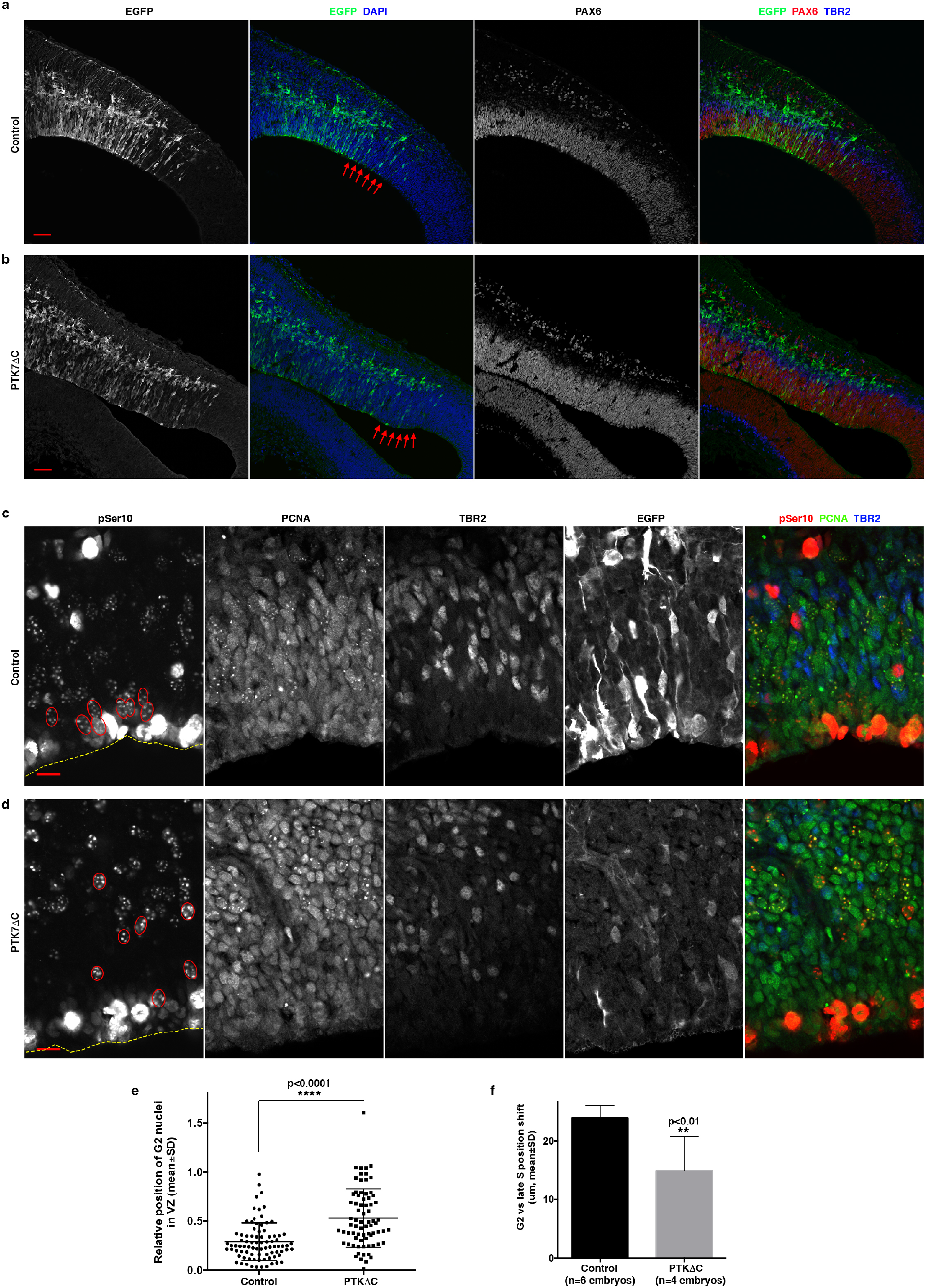
Expression of a mutant PTK7 results in increased thickness and convex ventricular surface in local neocortical areas. A plasmid for expressing PTK7ΔC was co-electroporated with an EGFP plasmid into the neocortex of wild-type mouse embryos at E12.5, and electroporated embryos were sacrificed 24h later. **a, b,** Expression of PTK7ΔC led to localized shape changes in the neocortex. Arrows in **b** indicate an area of increased thickness of the neocortex accompanied with convex ventricular surface in a PTK7ΔC transfected embryo. No such changes were observed in control transfected embryos (EGFP plasmid only) (arrows in **a**). Images are representative of 4 or more embryos for each group. Scale bars: 50μm. **c-f,** IKNM defects in PTK7ΔC transfected neocortices. Representative images are shown in **c** and **d**. Transfected neocortical areas with increased thickness were examined for IKNM defects. G2 nuclei (large puncta of pSer10 and diffuse nuclear PCNA) are outlined in red, and the ventricular surface is outlined by dashed yellow line. Note that many transfected cells (EGFP^+^) were poorly visualized by EGFP immunoreactivity because the samples were subjected to harsh antigen retrieval treatment -- which resulted in a weak to moderate EGFP signal as assessed by antibody staining. Scale bars: 10μm. **e**, The relative position of G2 phase NSC nuclei was significantly different between control and PTK7ΔC transfected neocortical areas. G2 phase NSCs pooled from 4 or more embryos for each group were used for quantification. **f,** The shift in position toward the ventricular surface in NSCs progressing from late S to G2 phase was significantly reduced in neocortical areas populated by transfected PTK7ΔC-expressing NSCs compared to control. **p<0.01, ****p<0.0001, Student’s t-test.

## PITP DKO neocortices exhibit cargo-specific ncPCP receptor membrane trafficking defects

PITPs potentiate Ptdns-4-P signaling-driven membrane trafficking from the TGN/endosomal system in yeast and in mammalian NSCs^5^. That body of evidence suggested that impairment of non-canonical Wnt/PCP signaling in PITP-deficient neocortex reflected a compromise in membrane trafficking of one or more ncPCP receptors from TGN/endosomes in PITP DKO NSCs. Indeed, pronounced accumulation of a punctate intracellular pool of VANGL2 was recorded in PITP DKO NSCs by immunostaining (Fig. 4a, b), and the punctate compartment of accumulation colocalized with the TGN marker Golgin97 (Fig. 4b). VANGL2 antibody specificity was verified by the nearly complete loss of immunoreactivity in neocortex expressing the unstable *lp* mutant form of VANGL2 (Extended Data Fig. 8)^18^. This TGN/endosomal membrane trafficking defect was also evident in PITP DKO embryos for VANGL1 (Extended Data Fig. 9a). By contrast, no such intracellular pools of PTK7 were detected in PITP DKO neocortex (Extended Data Fig. 9b), and the subcellular distribution of FZD2 (a receptor shared by the canonical Wnt and noncanonical Wnt/PCP signaling pathways) was also not obviously perturbed in PITP-deficient NSCs (Extended Data Fig. 9c). In aggregate, these data report that membrane trafficking of a subset of ncPCP receptors from the late Golgi system was significantly impaired in PITP DKO neocortices.

**Fig. 4.**
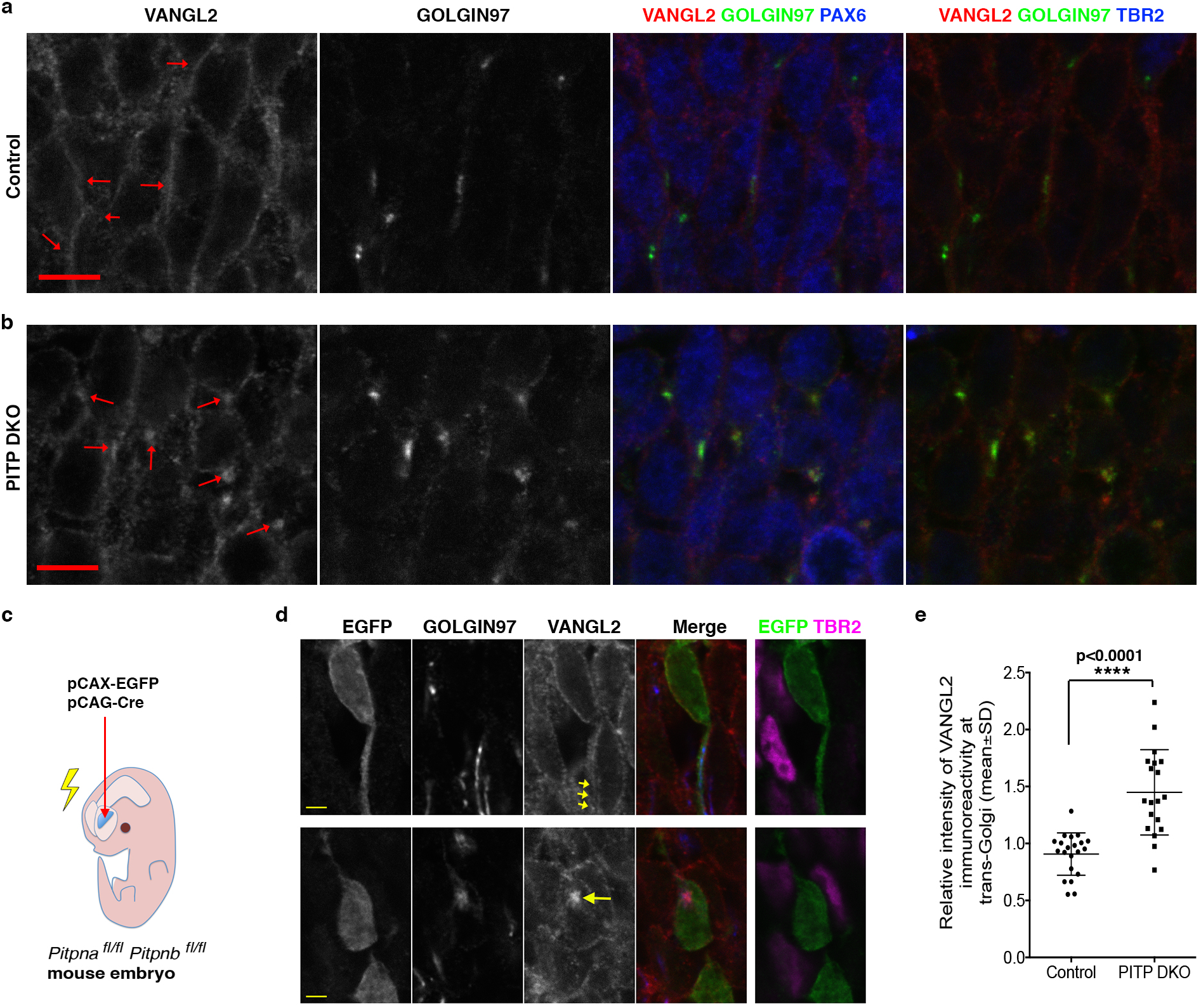
PITP-deficiencies lead to membrane trafficking defects of VANGL2. **a, b,** Accumulation of VANGL2 at trans-Golgi was detected in NSCs (PAX6^+^TBR2^−^) throughout the medial neocortex of PITP DKO neocortices at E12.5. Arrows indicate VANGL2 immunoreactivity at the NSC TGN (marked by GOLGIN97 labeling) in NSCs. VANGL2 immunoreactivity exhibits extensive co-localization with GOLGIN97 in PITP DKO neocortices but not in control neocortices. **c-e**, Deleting *Pitpna* and *Pitpnb* in individual NSCs in otherwise normal neocortices leads to accumulation of VANGL2 at trans-Golgi. **c**, schematic for the experiment. Control (*Pipna^+/+^ Pitpnb^+/+^*) or *Pipna^fl/fl^ Pitpnb^fl/fl^* mouse embryos were co-electroporated with an EGFP plasmid and a Cre plasmid at E12.5, and sacrificed at E16.5. Cre expression genetically ablated *Pitpna* and *Pitpnb* in transfected cells in *Pipna^fl/fl^ Pitpnb^fl/fl^* mouse embryos but not in control embryos. Non-transfected cells retain *Pitpna* and *Pitpnb* in both groups of embryos. **d**, Representative images. **e**, Quantification. VANGL2 immunoreactivity was significantly increased in TGN membranes of transfected NSCs (identified as TBR2^−^ cells located close to the ventricular surface) in *Pipna^fl/fl^ Pitpnb^fl/fl^* mouse embryos compared to control embryos. ****p<0.0001, Student’s t-test.

## ncPCP receptor membrane trafficking defects are cell-autonomous phenotypes and are GOLPH3-indepoendent

To determine whether VANGL2 membrane trafficking defects in PITP DKO neocortex represented cell-autonomous NSC phenotypes, and not indirect effects stemming from neocortical dysmorphologies in PITK DKO neocortices, the *Pitpna* and *Pitpnb* genes were ablated in a subset of NSCs residing in an otherwise wild-type neocortical niche. In these experiments, EGFP and Cre plasmids were co-introduced into E12.5 neocortices of *Pitpna^fl/fl^ Pitpnb^fl/fl^* embryos by in utero electroporation. Cre expression in individual transfected cells (marked with EGFP) induced deletion of both floxed *Pitpna* and *Pitpnb* alleles in those cells, whereas neighboring non-transfected cells remained competent for both *Pitpna* and *Pitpnb* gene expression^6^. As was recorded for PITP DKO neocortex, intracellular pools of VANGL2 accumulated in Golgin97-marked TGN/endosomal compartments of transfected NSCs. Such accumulations were not evident in adjacent untransfected NSCs (Fig. 4c-e). Thus, PITP DKO resulted in a cell-autonomous defect in VANGL2 trafficking from the NSC TGN/endosomal system.

PITP-dependent PtdIns-4-P synthesis on the TGN promotes anterograde cargo trafficking by stimulating phosphoinositide signaling pathways that enable efficient recruitment of PtdIns-4-P binding proteins/effectors to TGN membranes. One candidate for such an effector was GOLPH3 -- a peripheral membrane protein of TGN/endosomes whose role in promoting vesicle budding from the TGN is mediated via its interaction with the non-conventional actin-based motor Myo18A^24^. Moreover, GOLPH3 recruitment to NSC TGN membranes requires PITP-dependent PtdIns-4-P signaling, and this localization is essential for maintenance of NSC polarity in the developing neocortex^6^. Surprisingly, using an in utero electroporation regime analogous to the one described above, we found that functional silencing of GOLPH3 in transfected NSCs did not result in a detectable impairment of VANGL2 trafficking from TGN/endosomal membranes. That is, GOLPH3 deficits did not recapitulate the VANGL2 trafficking defects demonstrated by PITP DKO NSCs (Extended Data Fig. 10). These data indicate that, while PITPs potentiate VANGL2 trafficking from TGN/endosomes in a cell-autonomous manner, these do so via a mechanism that exhibits cargo-specificity and is substantially independent of GOLPH3 activity.

## Discussion

The mammalian neocortex is constructed via a highly ordered staging of neurogenesis and neuronal migration, and the tightly controlled spatial/temporal features of this developmental staging program are orchestrated by organization of the VZ. Consequently, regulation of VZ expansion in lateral vs radial dimensions plays a determining role in morphogenesis of the neocortex. Herein, we describe a critical role for the PITP/ncPCP axis in promoting IKNM in NSCs, and thereby regulating neocortical morphogenesis, via trafficking control of a subset of ncPCP receptors from the TGN/endosomal system. The data support the concept that IKNM is an essential feature of a specialized form of convergent extension that plays a fundamental role in driving the lateral expansion of the VZ at the expense of VZ expansion in the radial dimension.

In both yeast and mammalian NSCs, PITPs regulate membrane trafficking through the TGN/endosomal system by coordinating PtdIns-4-P signaling with PtdCho metabolism^5^ (our unpublished data). We therefore interpret the cell-autonomous VANGL1/2 trafficking defects recorded for PITP DKO NSCs as reflective of PtdIns-4-P signaling defects in the TGN/endosomal system of PITP-deficient NSCs. In that regard, the two unexpected findings encountered in this study were the cargo- and effector-specificities of these defects. That trafficking of only a subset of ncPCP receptors was compromised, while consistent with the viability of the PITP-deficient NSCs, nonetheless argues against models that presume all exocytic cargo is exported from the TGN/endosomal system in a PtdIns-4-P dependent manner^25^. An alternative interpretation is different TGN/endosomal cargo exhibit very different threshold requirements for PtdIns-4-P signaling in order to be efficiently exported from the TGN/endosomal system. The apparent PtdIns-4-P effector specificity was also unexpected. GOLPH3 is a PtdIns-4-P effector proposed to be a core component of the TGN trafficking machinery^24^. However, GOLPH3-deficient NSCs are viable and efficiently traffic VANGL2. We interpret these data to indicate other PtdIns-4-P effectors are primarily responsible for VANGL2 trafficking from the TGN/endosomal system to the plasma membrane. The cargo- and effector-specificities reported herein argue for functionally distinct PtdIns-4-P pools in the TGN/endosomal system rather than some bulk pool commonly accessed by PtdIns-4-P effectors. The ideas of PtdIns-4-P pool diversity and effector specificity converge conceptually as it suggests a mechanism where distinct PtdIns-4-P effectors engage with dedicated PtdIns-4-P pools to engineer what is in effect point resolution to PtdIns-4-P signaling on TGN/endosomal membrane surfaces^5^.

How does PITP/ncPCP signaling promote IKNM? An attractive possibility is that the PITP/ncPCP pathway couples cell adhesion signaling inputs at the NSC somata with modulation of actin cytoskeleton dynamics in the apical processes of NSCs. These actin dynamics then contribute to the forces that drive IKNM^3, 4^. Compromised trafficking to the plasma membrane of receptors of the ncPCP pathway is expected to diminish these cytoskeleton-based forces and result in defective IKNM. In that regard, it remains unclear why IKNM defects and neocortical shape changes were most pronounced in the medial neocortex of PITP DKO, *Vangl2^lp/lp^* and *Celsr1^ctb/ctb^* mutant embryos relative to the lateral neocortex. One formal possibility is that these phenotypes reflect regionally-specific mechanisms for promoting IKNM or regionally-specific IKNM functions. The existence of an unappreciated regional specificity in mechanisms of functional compensation in the lateral vs medial neocortex of PITP DKO, *Vangle2^lp/lp^* and *Celsr1^ctb/ctb^* embryos also remains a viable alternative interpretation of the data.

Finally, the results reported herein suggest interesting new possibilities when viewed in the context of autism spectrum disease. It has long been appreciated that fluctuations in brain inositol levels are correlated with pre-dementia, mood disorders and cognitive impairments in humans^26^. Indeed, there is growing support for inositol supplementation as a nutraceutical approach for preventing and treating cognitive insufficiencies and other neurodegenerative disorders^26^. In that regard, we are intrigued by the fact that human *PTK7* is a high-confidence autism-risk gene^27^. One implication of our data is that deranged PITP/ncPCP signaling, and the associated deficits in IKNM and neocortical morphogenesis, integrate these seemingly disparate sets of clinical observation. That is, defective PITP/ncPCP signaling contributes to structural and functional deficits in the neocortex that expose the developing fetus to elevated autism-risk.

## Acknowledgements

We thank C. S. Abrams for providing the *Pitpna*-floxed and *Pitpnb*-floxed mice. This work was supported by NIH grant R35 GM131804 (to V.A.B.) and grant B-0017 from the Robert A. Welch Foundation (to V.A.B.)

## Author contributions

Z.X. and V.A.B. designed the experiments. Z.X. performed all experiments. Z.X. and V.A.B. wrote the manuscript.

## METHODS

### Plasmids

Plasmids for expression of EGFP (pCAX-EGFP), Cre (pCAG-Cre), and *Golph3* shRNA were previously described^6^. To generate plasmids for the expression of PTK7ΔC, the coding sequence for PTK7 amino acid 1-728 (lacking the cytoplasmic domain) was amplified via RT-PCR from neocortex RNA preparations of E14.5 mouse embryos, and then inserted into the pCAX vector. RT-PCR was performed using iScript Select cDNA synthesis Kit (BIO-RAD) and high-fidelity Phusion DNA polymerase (New England Biolabs). The plasmid was verified by DNA sequencing.

### Antibodies

The antibodies used include: rabbit polyclonal anti-GOLGIN97 (Abcam, ab98023), chicken polyclonal anti-GFP (Aves Labs, GFP-1010), mouse monoclonal anti-PAX6 (Developmental Studies Hybridoma Bank, PAX6), rabbit polyclonal anti-PAX6 (LSBio, LS-C179903 and LS-B16455), rabbit polyclonal anti-TBR2 (Abcam, ab183991), rat polyclonal anti-TBR2 (Thermo Fisher Scientific, 14-4875-82), sheep polyclonal anti-VANGL2 (Novus Biologicals, AF4815-SP), rabbit polyclonal anti-VANGL1 (Novus Biologicals, NBP1-86990), rabbit polyclonal anti-PTK7 (ProteinTech Group, 17799-1-AP), goat polyclonal anti-FZD2 (Novus Biologicals, NBP1-20920), goat polyclonal anti-PCNA (LSBio, B14132), and rabbit polyclonal anti-pSer10 (GenScript, A00339). Secondary antibodies (with minimal species cross reactivity) were from Jackson Immuno Research Laboratories, Inc (West Grove, PA).

### Mice and in utero electroporation

The *Pitpna*/*Pitpnb*-floxed mice and *Emx1*-Cre mice were on the C57BL6 background and described previously^6^. The *Celsr1^ctb^* mice and the *Vangl2^lp^* mice were obtained from The Jackson Laboratory (*Celsr1^ctb^*: stock#016111, BALB/cByJ background; *Vangl2^lp^*: stock#000220, mixed C57BL/6J and C3H background). Mice were maintained in the animal facility of Texas A&M University Health Science Center and handled in accordance with National Institute of Health and institutional guidelines on the care and use of animals. The gender of mouse embryos used in these experiments was not determined. For embryonic staging, the day of vaginal plug detection was designated as E0.5. In utero electroporation was performed as previously described^6^. Briefly, timed-pregnant female mice were anesthetized and subjected to laparotomy to expose the uteri. The brain of mouse embryos was visualized using a fiber optic light source. Plasmid solutions were then injected into the lateral ventricle of the embryonic mouse brain through the uterine wall. Following plasmid injection, 5 short electric pulses were delivered across the head of mouse embryos. The uteri were then returned to the abdominal cavity of the dam, and the incision was sutured. The dam was allowed to recover at a warm location before being transferred to animal holding room. For *Pitpna*/*Pitpnb* knockout electroporation experiments, a mixture of pCAX-EGFP and pCAG-Cre (mass ratio of 1:2) was introduced into the neocortex of mouse embryos (*Pitpna^fl/fl^ Pitpnb^fl/fl^* or *Pitpna^+/+^ Pitpnb^+/+^*) at E12.5, and electroporated embryos were harvested at E16.5. For GOLPH3 knockdown electroporation experiments, a mixture of pCAX-EGFP and Golph3 shRNA (mass ratio of 1:3) was introduced into the neocortex of wild-type mouse embryos at E12.5, and electroporated embryos were harvested a E15.5. For PTK7ΔC electroporation experiments, a mixture of pCAX-EGFP and pCAX-PTK7ΔC (mass ratio of 1:5) was introduced into the neocortex of wild-type mouse embryos at E12.5, and electroporated embryos were harvested 24h later.

### Genotyping

Genomic DNA from mouse ear punching tissues was used for PCR genotyping. Taq DNA polymerase was used for genotyping PCR reactions. Genotyping for floxed *Pitpna*, floxed *Pitpnb*, and *Emx1-Cre* alleles was previously described^6^. The following primers were used for genotyping the *ctb* line of *Celsr1* mice: Celsr1del1-F2wt (GTTCCTGGAGGTGTGAGC); Celsr1del1-F2del1 (GTTCCTGGAGGTGTGAGG); Celsr1del1-R2 (GAGCAGCTCCTGGAATCTTTG). The pair of Celsr1del1-F2wt and Celsr1del1-R2 detects the wild-type *Celsr1* allele (300bp; annealing temperature at 60°C), whereas the pair of Celsr1del1-F2del1 and Celsr1del1-R2 detects the mutant *Celsr1^ctb^* allele (300bp; annealing temperature at 60°C). For genotyping the *lp* line of *Vangl2* mice, a PCR product of 241bp encompassing the location of the *lp* mutation was amplified from the genomic DNA preparations using primers LpSeq-F (GAAAGCTGTGTCACAGAGGATG) and LpSeq-R (TCACTACCAAGCTGAAGTCCTG). The genotypes were determined by sequencing analysis of this PCR product.

### Tissue preparation and immunostaining

For tissue sample preparation of E12.5 embryos, the whole brain was dissected out in PBS and fixed in 2% paraformaldehyde (in 1X PBS) for 20-30min. For tissue sample preparation of older embryos, forebrain hemispheres were dissected out, part of the hippocampus was removed to expose the lateral ventricle, and the rest of the forebrain was fixed in 2% paraformaldehyde (in 1X PBS) for 25-30min. Fixed forebrain samples were cryoprotected with 20% sucrose (in 1X PBS), embedded in Tissue-Tek OCT, and cryosectioned at the thickness of 20-30μm. Sections were stored at −20°C before the immunostaining procedure. For immunostaining, both primary and secondary antibodies were diluted in 1xPBS containing 3% bovine serum albumin and 0.2% Triton-X-100.

Antibody incubation steps (primary antibody: overnight; secondary antibody: 1-3h) were performed in a humidified chamber protected from direct light. Secondary antibodies with the following cyanine dyes were used: Cy2 (green), Cy3 (red), Cy5 (far red), and DyLight405 (blue). EverBrite™ mounting media from Biotium (Fremont, CA) were used to mount coverslips and protect from photo bleaching.

### Antigen retrieval treatment

Brain sections were subjected to antigen retrieval treatment prior to PCNA and pSer10 immunostaining. The procedure was performed similarly as previously described^28^. Briefly, tissue sections were submerged in 1x antigen retrieval citrate solution (Sigma), microwaved for 5 min on power 90, left in the heated buffer for 5 min, and washed in 1x PBS three times prior to the immunostaining procedure.

### Confocal microscopy and immunoreactivity quantifications

Confocal images were obtained on a Nikon TiE confocal microscope using the NIS-Elements software. Quantification of relative immunoreactivity of VANGL2 was performed similarly as previously described for that of GOLPH3^6^. Briefly, Confocal images were converted to TIF files, which were then used for gray value measurement in Adobe Photoshop CS6. Cells from images obtained from at least three embryos were pooled together for each group. To quantify relative VANGL2 immunoreactivity at trans-Golgi for each transfected cell (EGFP^+^), VANGL2 immunoreactivity at trans-Golgi (indicated by GOLGIN97 immunoreactivity) was compared to the thin layer of VANGL2 immunoreactivity surrounding the nucleus, which was consistent with the pattern of plasma membrane localization at the somata of NSCs.

### Statistical Analysis

The GraphPad Prism software (version 6.0b) was used for statistical analysis. Unpaired, two-tailed Student’s t-test was used to compare results from two groups. One-way ANOVA were used to compare results from three or more groups.

### Ethical regulations

All experiments were performed in compliance with relevant ethical regulations. Mice were maintained in the animal facility of Texas A&M University Health Science Center and handled in accordance with National Institute of Health and institutional guidelines on the care and use of animals.

## Data availability

All data will be provided by the corresponding authors upon reasonable request.

---

**Extended Data Fig. 1.**
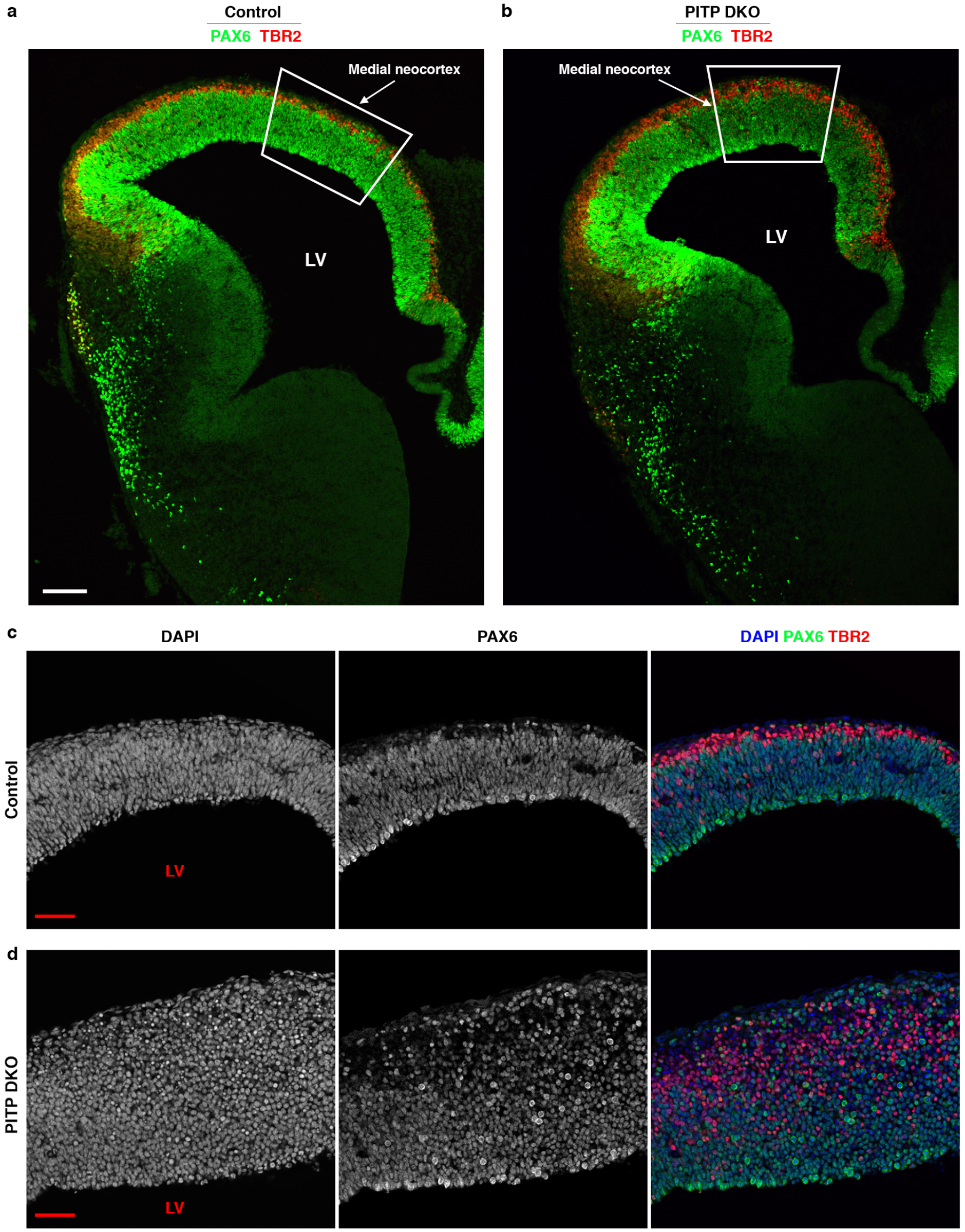
Shape and cell components in control and PITP-deficient neocortices. **a, b,** Shape of the neocortex at early E12. Compared to control (genotype: *Pitpna^fl/+^ Pitpnb^fl/fl^ Emx1^Cre/+^*), the medial neocortex (boxed area) in PITP DKO embryos (genotype: *Pitpna^fl/fl^ Pitpnb^fl/fl^ Emx1^Cre/+^*) was thicker radially and shorter mediolaterally. Images are representative of at least 4 embryos for each group. **c, d,** Most cells near the ventricular surface (which surrounds LV) were NSCs in both control and PITP DKO neocortices at late E12. NSCs and IPCs were identified as PAX6^+^TBR2^−^ cells and TBR2^+^ cells, respectively. Images are representative of at least 4 embryos for each group. LV: lateral ventricle. Scale bars: 100μm in **a**, **b**, 50μm in **c**, **d**.

**Extended Data Fig. 2.**
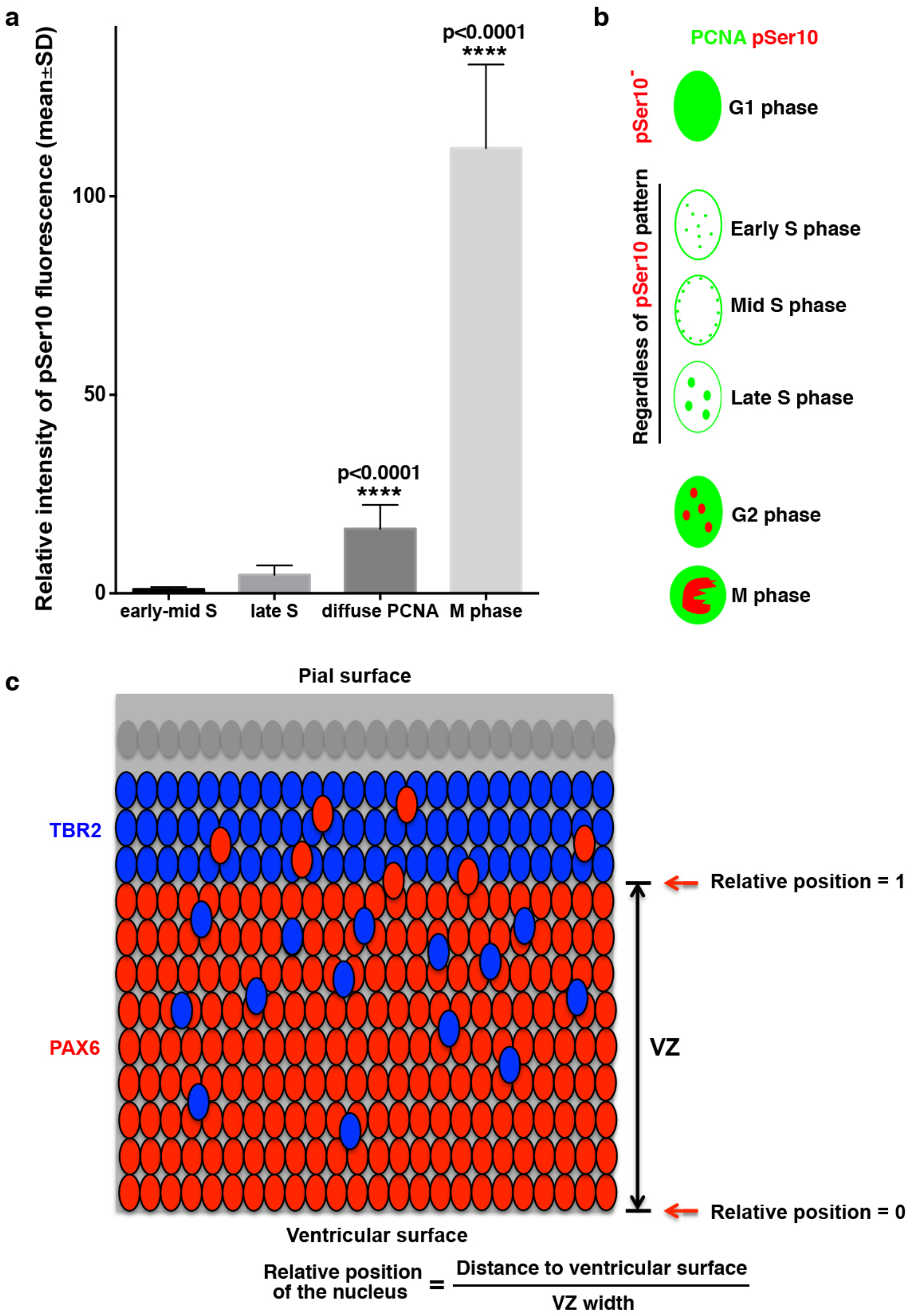
Analysis of cell cycle phases in the embryonic mouse neocortex. **a,** Intensity of pSer10 immunofluorescence in different subpopulations of NSCs at E12.5. pSer10 immunofluorescence was weakly detected in early to mid S phase NSCs (small nuclear puncta of PCNA), slightly increased in late S phase NSCs (large nuclear puncta of PCNA), further increased in pSer10^+^ NSCs with diffuse nuclear PCNA, and drastically increased in M phase NSCs (intense pSer10^+^ chromatin with cytoplasmic PCNA). **** p < 0.0001, one-way ANOVA. **b,** Identification of cell cycle phases based on fluorescence patterns of PCNA and pSer10. G1: pSer10^−^, diffuse nuclear PCNA; early S: small nuclear puncta of PCNA, mostly not at the nuclear periphery; mid S: small nuclear puncta of PCNA, mostly at the nuclear periphery; late S: large nuclear puncta of PCNA; G2: diffuse nuclear PCNA and large nuclear puncta of pSer10; M: cytoplasmic PCNA and intense pSer10^+^ chromatin. **c**, Schematic for nuclear position quantification. The upper border of VZ is delineated by the layer of densely packed IPCs (TBR2^+^).

**Extended Data Fig. 3.**
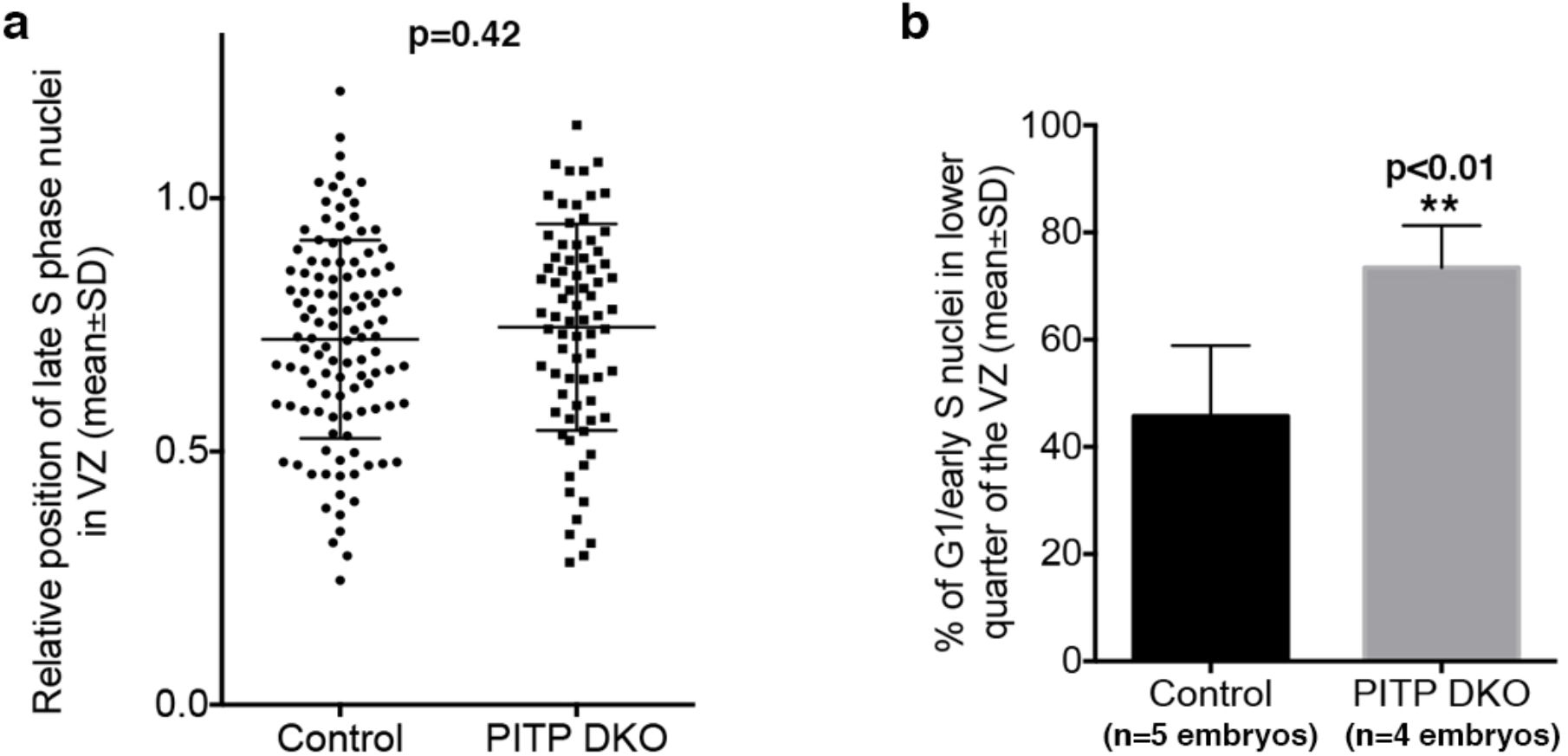
Analysis of IKNM defects in PITP-deficient neocortices at E12.5. **a,** Relative position of late S phase NSCs in PITP DKO neocortices was similar to that in control neocortices. NSC nuclei (PAX6^+^TBR2^−^) pooled from 4 different embryos for each group were quantified and analyzed by Student’s t-test. **b**, The percentage of G1 and early S phase NSC nuclei at the apical quartile of the VZ was increased in PITP DKO neocortices. **p<0.01, Student’s t-test.

**Extended Data Fig. 4.**
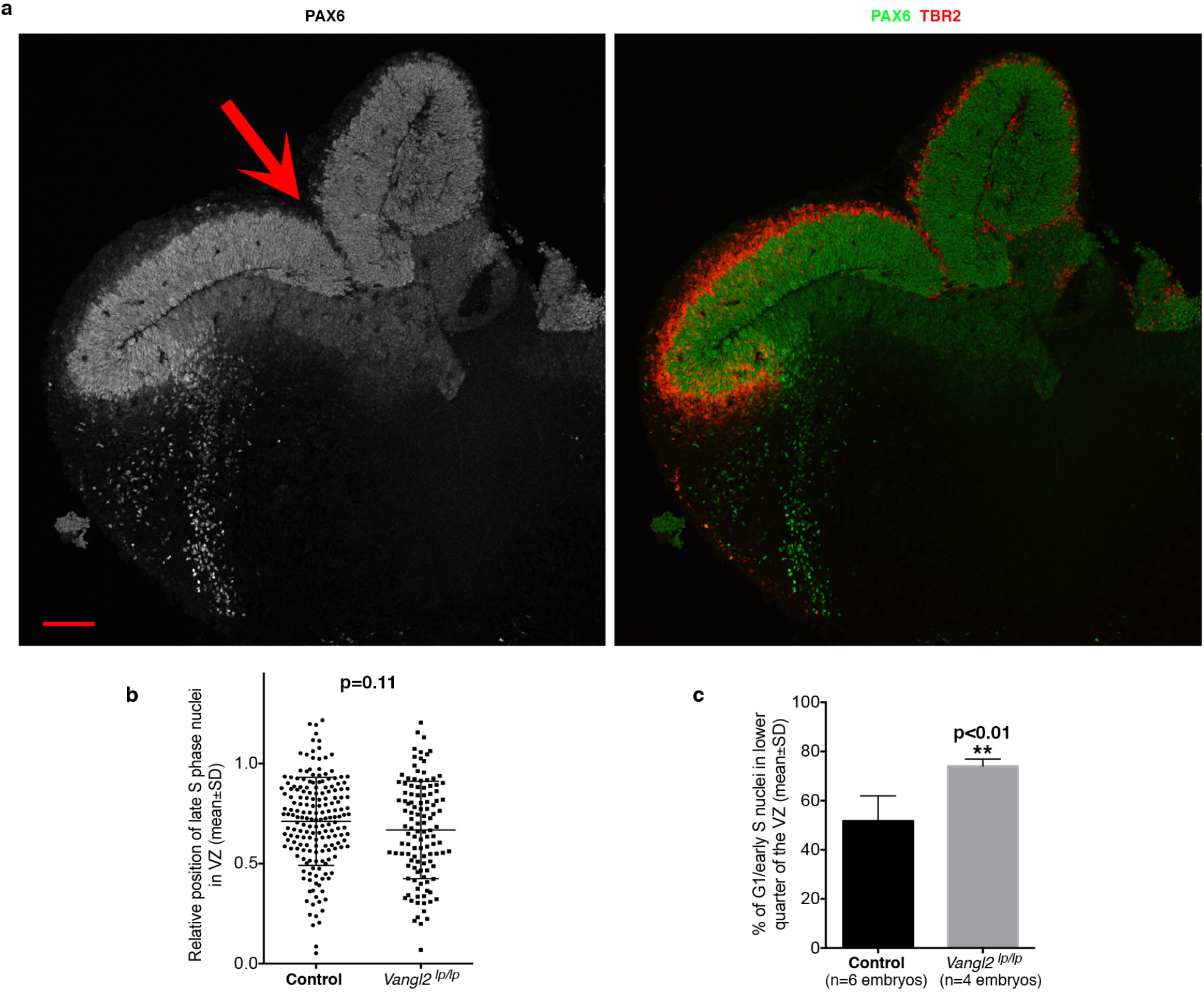
Neocortical shape and IKNM in *Vangl2^lp/lp^* mutant embryos at E12.5. **a,** The neocortex was typically broken at one location (indicated by the arrow) in *Vangl2^lp/lp^* mutant embryos. Images are representative of 5 embryos. Scale bar: 100 μm. **b,** The relative position of late S phase NSCs in the VZ was similar between *Vangl2^lp/lp^* mutant embryos and control (*Vangl2^+/+^* or *Vangl2^lp/+^*) embryos. NSCs pooled from at least 4 embryos for each group were quantified. **c,** The percentage of G1/early S phase NSC nuclei in the apical quartile of the VZ was increased in the neocortex of *Vangl2^lplp^* mutant embryos. **p<0.01, Student’s t-test. Neocortical areas with increased thickness were quantified in **b** and **c**.

**Extended Data Fig. 5.**
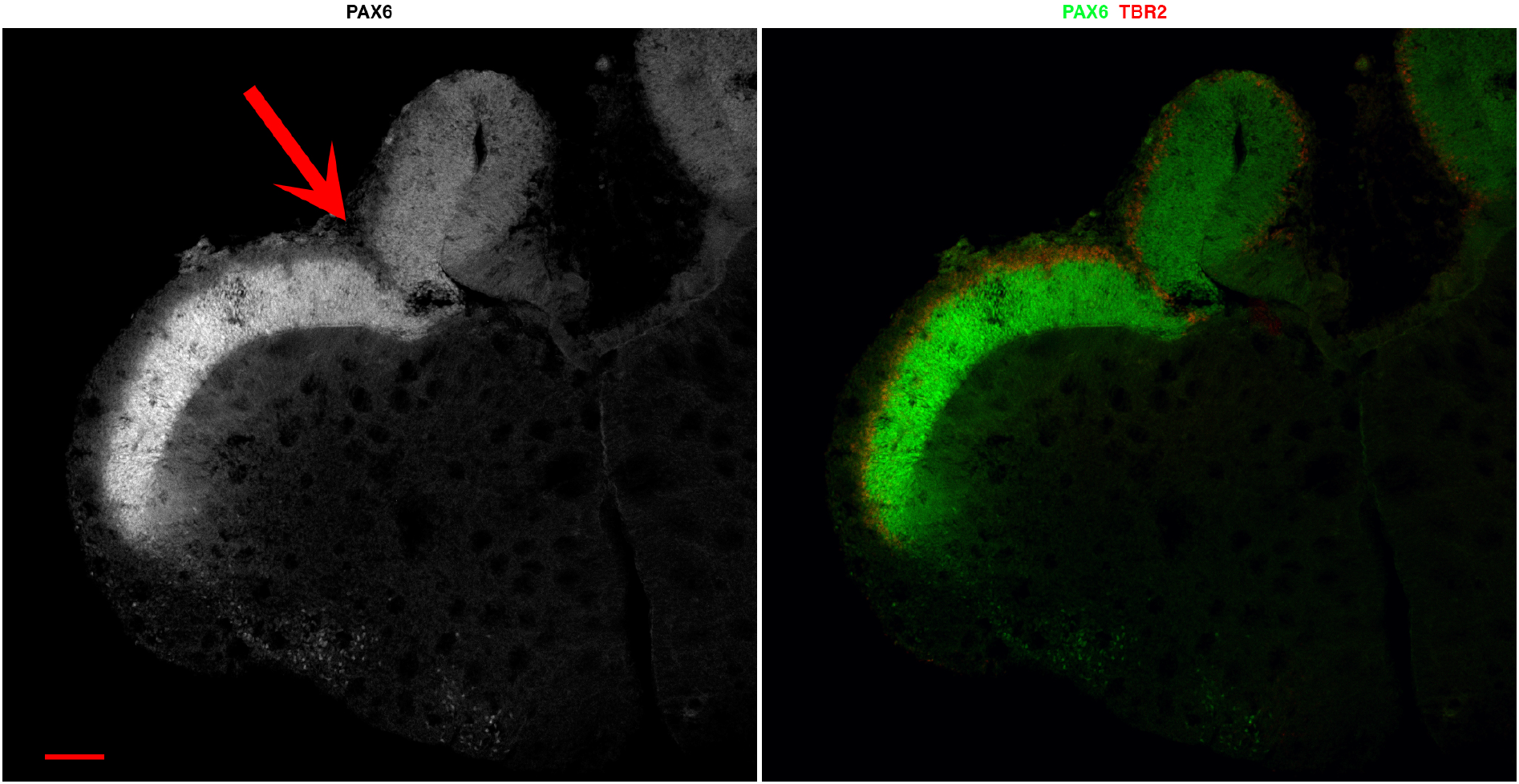
Broken neocortex in *Celsr1^ctb/ctb^* mutant embryos at E12.5. The arrow indicates the breaking point. Images are representative of 6 embryos. Scale bar: 100 μm.

**Extended Data Fig. 6.**
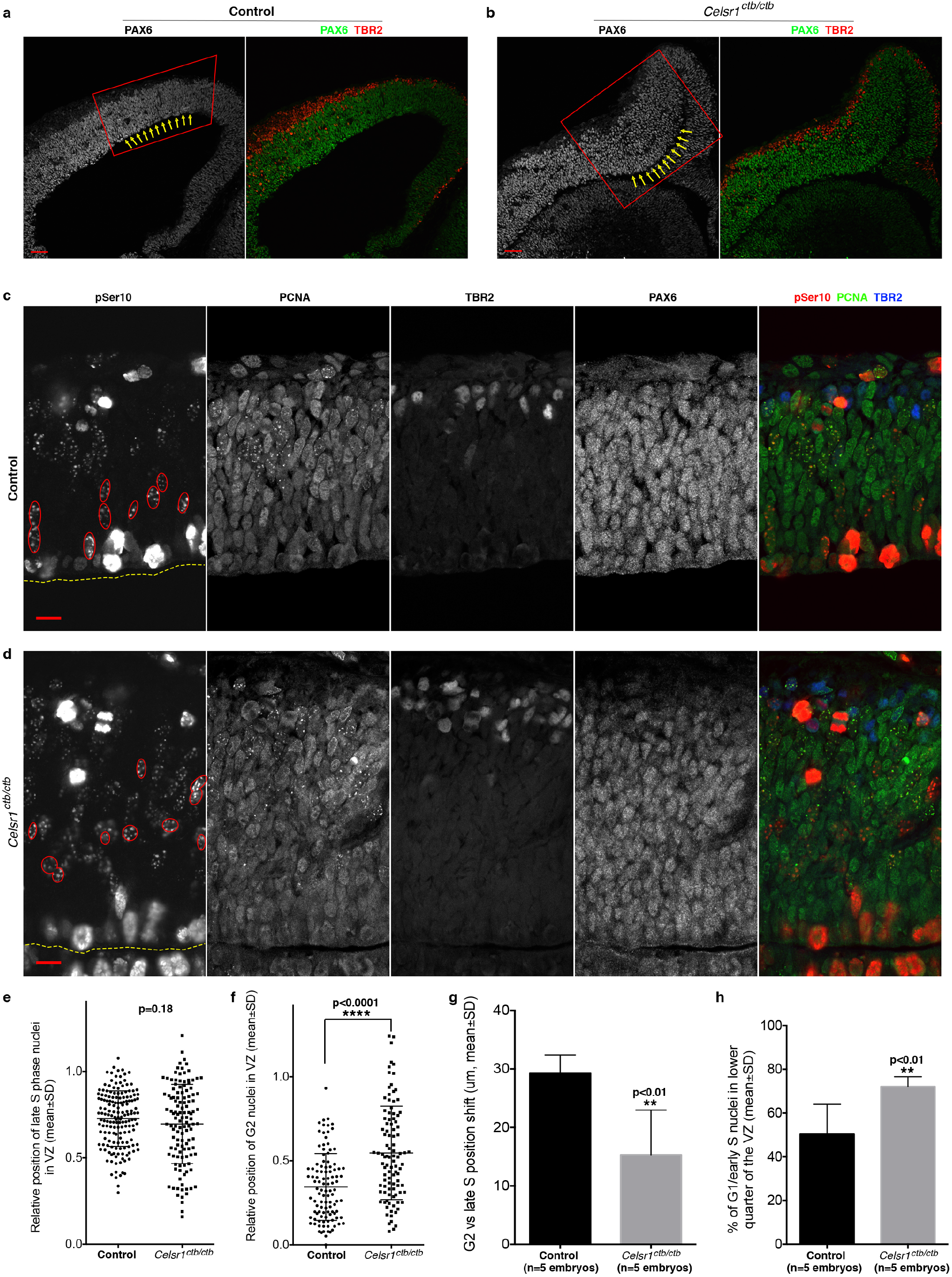
Altered neocortical shape and impaired IKNM in *Celsr1^ctb/ctb^* mutant embryos. **a, b,** Increased thickness (boxed areas) and convex ventricular surface (indicated by arrows in **b,** compared to the area indicated by arrows in **a**) were common in *Celsr1^ctb/ctb^* mutant embryos at E12.5, particularly in the medial part of the neocortex. Images are representative of at least 6 embryos for each group. Scale bars: 50μm. **c-h,** IKNM defects in *Celsr1^ctb/ctb^* mutant embryos at E12.5. Representative images are shown in **c** and **d**. Neocortical areas with increased thickness were examined for possible IKNM defects. G2 nuclei (large puncta of pSer10 and diffuse nuclear PCNA) are outlined in red, and the ventricular surface is outlined by dashed yellow lines. Scale bars: 10μm. Various quantifications of nuclear position are shown in **e-h**. **e**, The position of late S phase NSC nuclei was similar between control and *Celsr1^ctb/ctb^* mutant embryos. **f,** The relative position of G2 phase NSC nuclei was less shifted to the lower part of the VZ in *Celsr1^ctb/ctb^* mutant embryos compared to control. **g**, The overall change of position (shift toward the ventricular surface) from late S phase to G2 phase was significantly reduced in *Celsr1^ctb/ctb^* mutant embryos compared to control. **h**, The percentage of G1/early S phase NSC nuclei in the apical quartile of the VZ was significantly increased in *Celsr1^ctb/ctb^* mutant embryos compared to control. **p<0.01, ****p<0.0001, Student’s t-test was used for all analyses.

**Extended Data Fig. 7.**
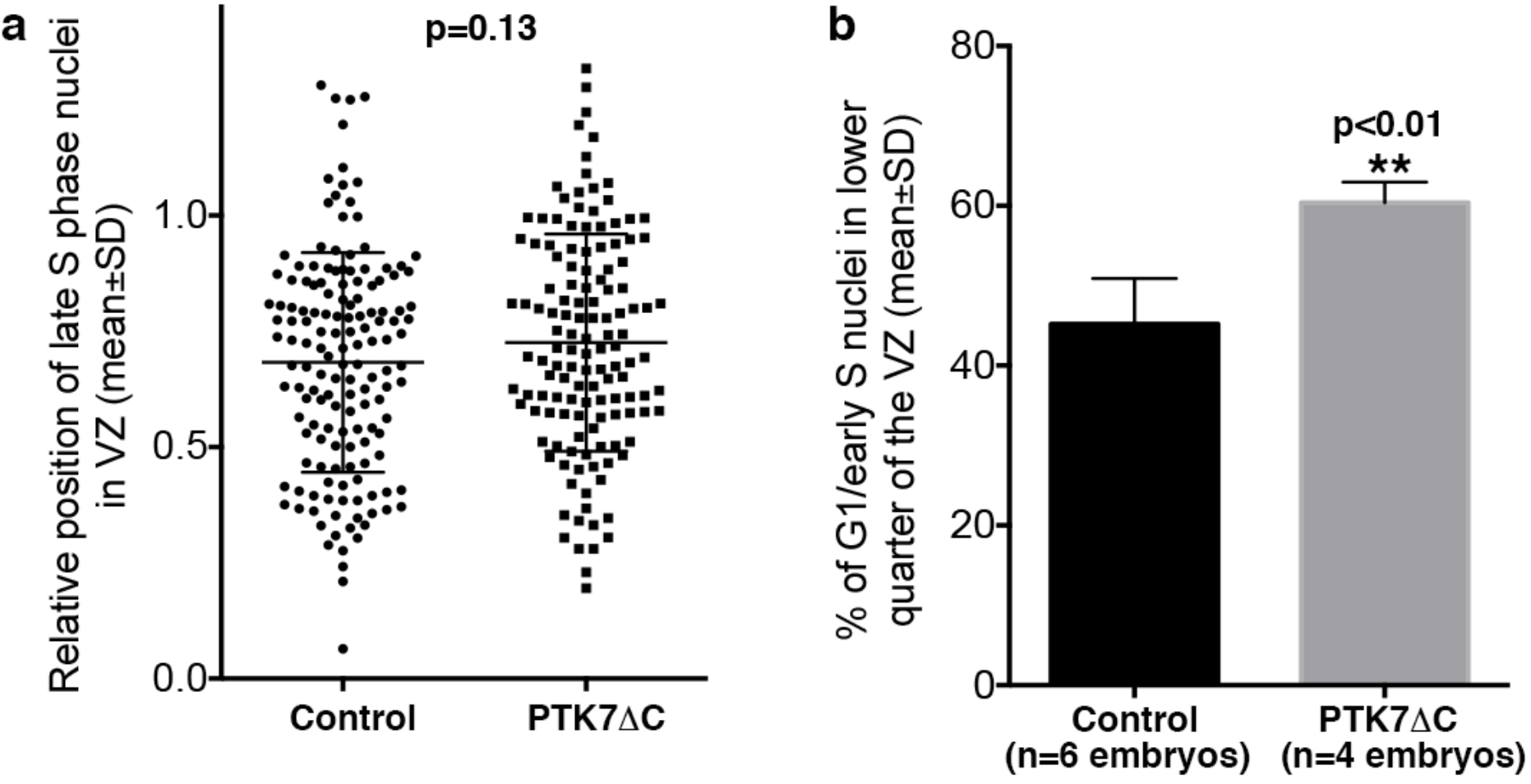
Analysis of IKNM defects in PTK7-deficient neocortices. Mouse embryos were electroporated at E12.5 and sacrificed for immunostaining analysis 24h later. **a,** Relative position of late S phase NSCs in PTK7ΔC-transfected neocortices was similar to that in control-transfected neocortices. NSC nuclei (PAX6^+^TBR2^−^) pooled from 4 or more different embryos for each group were quantified and analyzed by Student’s t-test. **b**, The percentage of G1 and early S phase NSC nuclei at the apical quartile of the VZ was increased in PTK7ΔC-transfected neocortices. **p<0.01, Student’s t-test.

**Extended Data Fig. 8.**
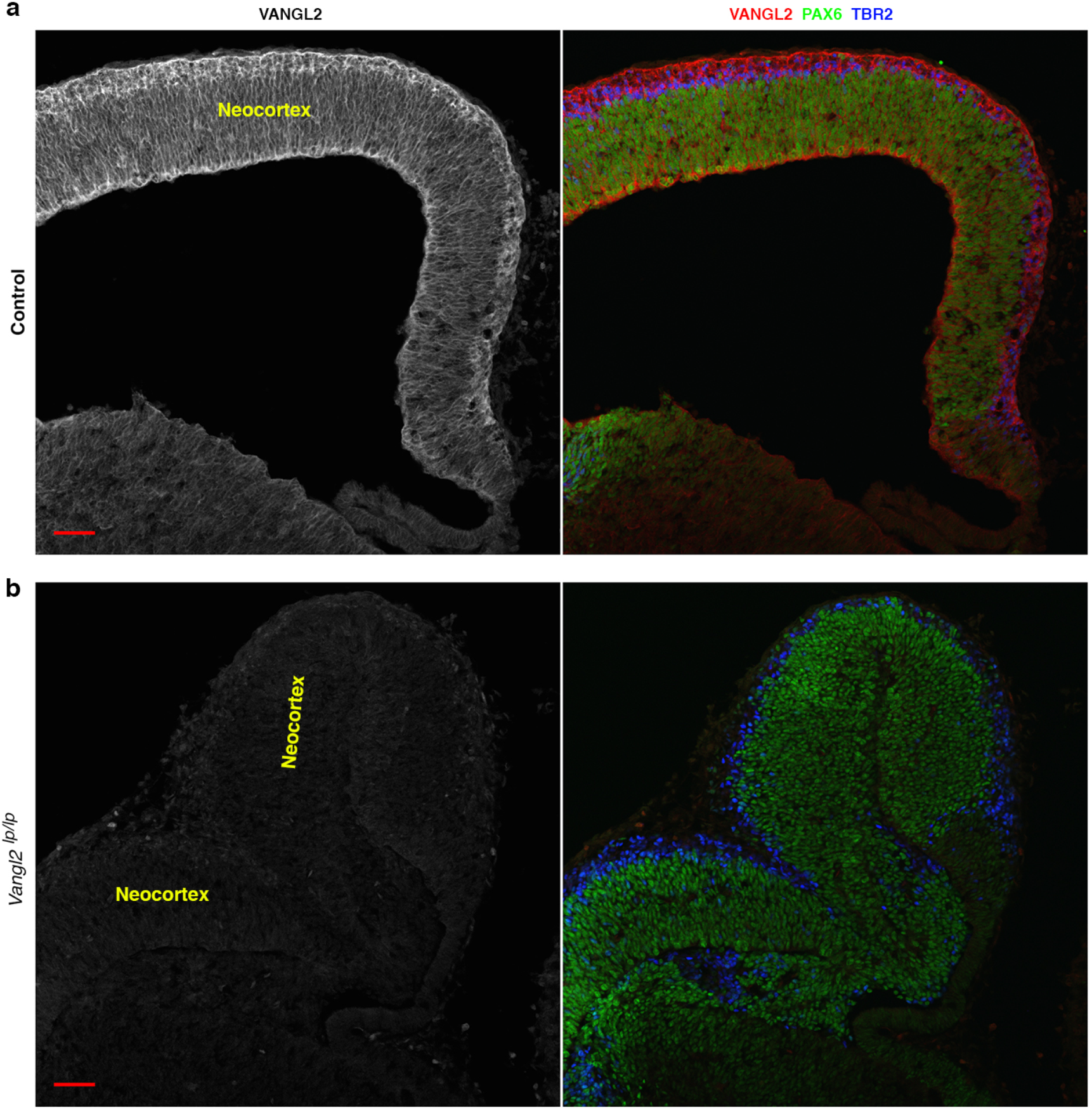
Specificity of the VANGL2 antibody used in this study. The neocortices of E12.5 mouse embryos were immunostained using a sheep polyclonal antibody against VANGL2. The immunofluorescence intensity of VANGL2 labeling was markedly reduced in the neocortices (and other brain areas) of *Vangl2^lplp^* embryos compared to that in control embryos. The *lp* mutant of VANGL2 protein is highly unstable, and therefore at markedly reduced levels than the wild-type VANGL2 protein^18^.

**Extended Data Fig. 9.**
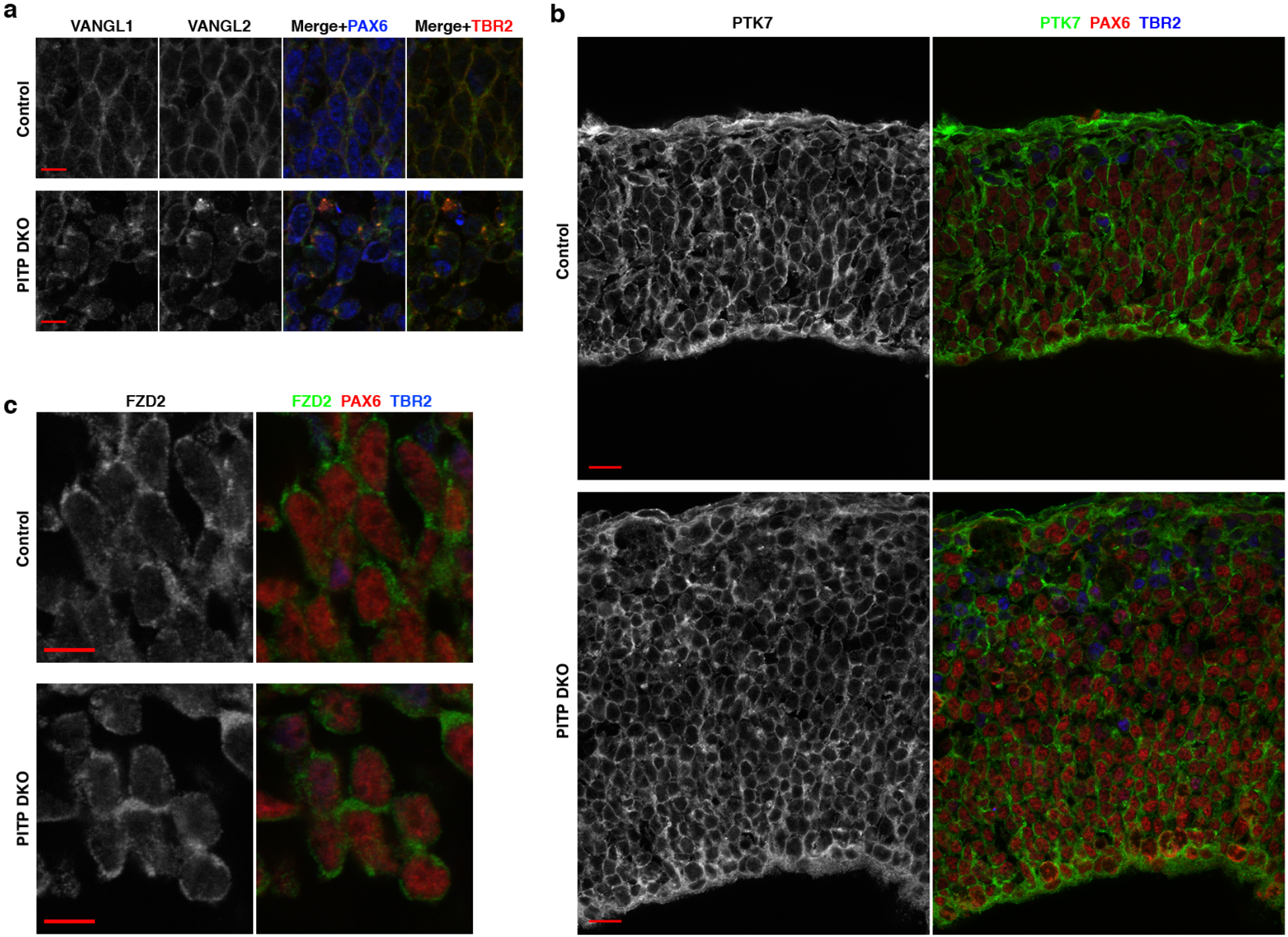
Subcellular distribution of various receptors in PITP DKO neocortices at E12.5. **a,** The pattern of VANGL1 immunoreactivity was similarly changed as that of VANGL2 immunoreactivity in PITP DKO neocortices. **b, c,** The patterns of PTK7 immunoreactivity and FZD2 immunoreactivity were largely the same between control and PITP DKO neocortices. Scale bars: 5μm in **a**, **c**, and 10μm in **b**.

**Extended Data Fig. 10.**
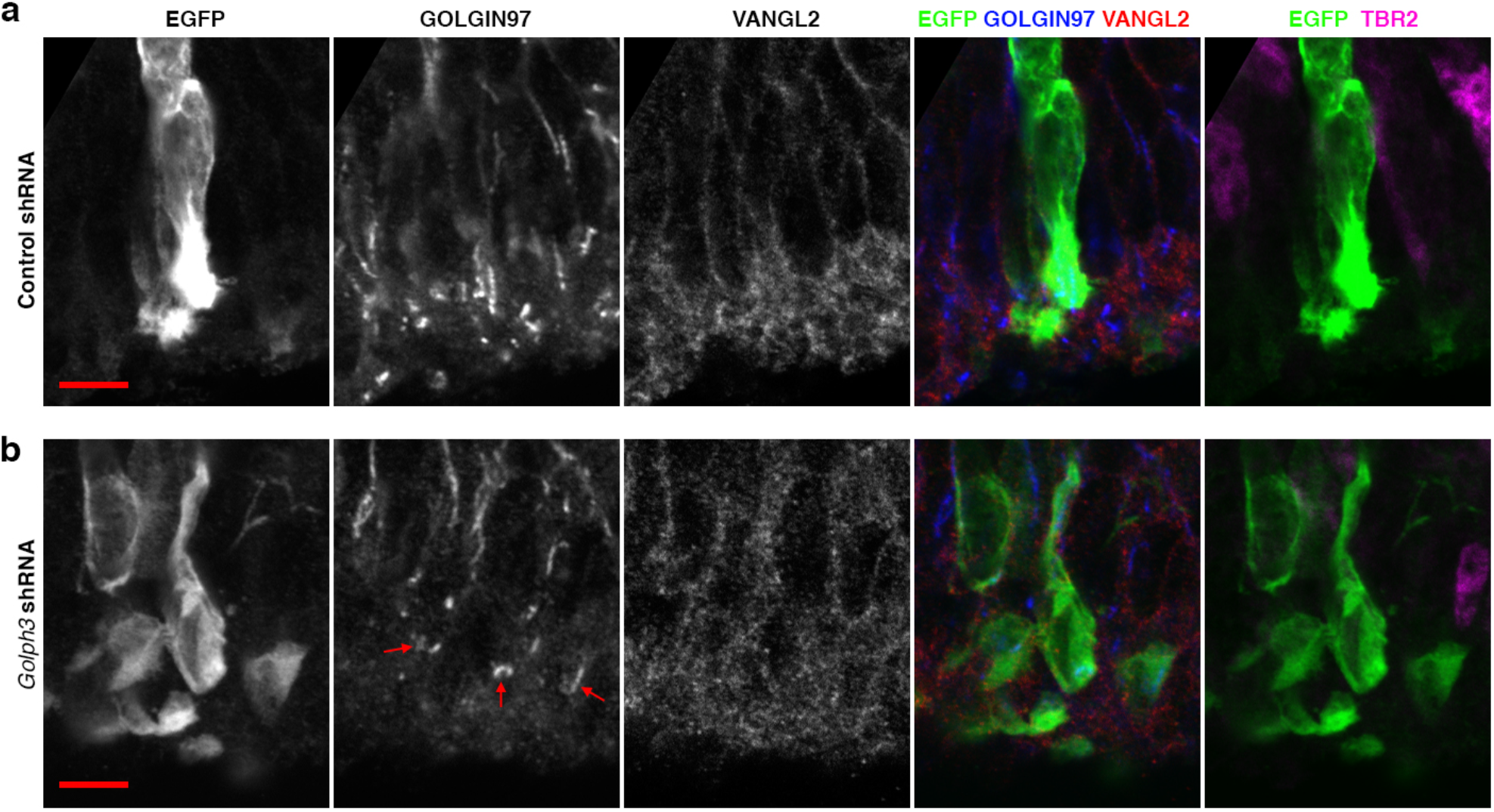
GOLPH3 knockdown does not cause perinuclear accumulation of VANGL2 in the embryonic mouse neocortex. Control or *Golph3* shRNA^6^ was introduced into the neocortex of mouse embryos at E12.5 via in utero electroporation, and electroporated embryos were sacrificed at E15.5. VANGL2 immunoreactivity in transfected NSCs (EGFP^+^TBR2^−^ cells located close to the ventricular surface) was similar between control (**a**) and the *Golph3* shRNA group (**b**). Images are representative of results from three or more embryos for each group. Arrows in **b** point to the trans-Golgi network (identified by GOLGIN97 immunoreactivity) in several transfected NSCs. Scale bars: 5μm.

